## Supplementary data Figures 2-4 for "NGS method for parallel processing of high quality, damaged or fragmented input material using target enrichment"

#### FIGURE 2

| | Concentration (ng/ul) | | | |
|---|---|---|---|---|
| Sample name | HyperPrep | HyperPlus 6.5 min | HyperPlus 12.5 min | HyperPrep + HyperPlus 12.5 min |
| FFPE-1 | 25.9 | 11.7 | 14.5 | 26.3 |
| FFPE-2 | 38.2 | 35.4 | 41.9 | 59.1 |
| FFPE-3 | 53.4 | 35.3 | 39.7 | 56.8 |
| FFPE-4 | 34.3 | 75.0 | 100.3 | 113.1 |
| NA24143 | 14.8 | 227.0 | 259.0 | 277.8 |

#### FIGURE 3

| A | Mean insert size (bp) | | |
|---|---|---|---|
| Sample name | HyperPrep | HyperPlus | HyperPrep + HyperPlus |
| FFPE-1 | 118 | 110 | 110 |
| FFPE-2 | 159 | 136 | 140 |
| FFPE-3 | 137 | 125 | 125 |
| FFPE-4 | 243 | 161 | 166 |
| NA24143 | - | 224 | 224 |

| B | Duplicates | | |
|---|---|---|---|
| Sample name | HyperPrep | HyperPlus | HyperPrep + HyperPlus |
| FFPE-1 | 0.69 | 0.69 | 0.64 |
| FFPE-2 | 0.59 | 0.51 | 0.49 |
| FFPE-3 | 0.49 | 0.49 | 0.47 |
| FFPE-4 | 0.57 | 0.33 | 0.34 |
| NA24143 | - | 0.25 | 0.28 |

| C | Median target coverage (X) | | |
|---|---|---|---|
| Sample name | HyperPrep | HyperPlus | HyperPrep + HyperPlus |
| FFPE-1 | 121 | 78 | 130 |
| FFPE-2 | 124 | 160 | 201 |
| FFPE-3 | 80 | 58 | 105 |
| FFPE-4 | 251 | 406 | 405 |
| NA24143 | - | 976 | 955 |

| D | PCT_TARGET_BASES_150X | | |
|---|---|---|---|
| Sample name | HyperPrep | HyperPlus | HyperPrep + HyperPlus |
| FFPE-1 | 0.346 | 0.262 | 0.419 |
| FFPE-2 | 0.440 | 0.518 | 0.621 |
| FFPE-3 | 0.421 | 0.410 | 0.444 |
| FFPE-4 | 0.569 | 0.637 | 0.636 |
| NA24143 | - | 0.998 | 0.998 |

### FIGURE 4

| Sample name | PCT_TARGET_BASES_500X | PCT_TARGET_BASES_1000X | FRACTION_DUPLICATES | MEAN_INSERT_SIZE | MEDIAN_TARGET_COVERAGE |
|---|---|---|---|---|---|
| NA24143-10ng | 0.947 | 0.002 | 0.657 | 223 | 720 |
| NA24143-50ng | 0.998 | 0.977 | 0.262 | 224 | 1782 |
| NA24143-250 ng | 0.998 | 0.991 | 0.194 | 254 | 2001 |
| HD798-50 ng | 0.994 | 0.853 | 0.299 | 220 | 1609 |
| HD798-250 ng | 0.998 | 0.984 | 0.204 | 209 | 1921 |
| HD799-50 ng | 0.988 | 0.768 | 0.426 | 170 | 1239 |
| HD799-250 ng | 0.997 | 0.965 | 0.241 | 168 | 1734 |
| HD803-50 ng | 0.798 | 0.349 | 0.538 | 130 | 847 |
| HD803-250 ng | 0.967 | 0.774 | 0.305 | 127 | 1367 |
| HD777-10 ng | 0.140 | 0.000 | 0.776 | 191 | 429 |
| HD777-30 ng | 0.996 | 0.800 | 0.470 | 184 | 1207 |
